# Transforming dairy waste into hydrogen fuel using alginate-encapsulated bacterial co-cultures

**DOI:** 10.64898/2025.12.18.695156

**Authors:** Danielle T. Bennett, Ryan M. Kosko, Farwa Awan, Todd D. Krauss, Kara L. Bren, Anne S. Meyer

## Abstract

Dairy waste, such as whey resulting from cheese production, is produced in massive volumes worldwide and is regarded as environmentally difficult to dispose of due to its high organic content^1^. Harnessing the potential of this waste material to support the synthesis of valuable products such as hydrogen fuel or reduced graphene oxide, which may be utilized for conductive thin films and energy storage^2,3^, can reduce waste and add revenue streams for dairy farmers. Here, we demonstrate a circular bioeconomy using alginate-encapsulated co-cultures of *Shewanella oneidensis* together with lactic-acid-producing bacteria *Klebsiella pneumoniae*. These co-cultures can directly metabolize unprocessed cheese-making waste as an electron source instead of costly, environmentally high-impact lactic acid^4,5^. Alginate-encapsulated co-cultures fed unprocessed dairy waste showed a 2-to-3-fold higher graphene oxide reduction rate compared to *S. oneidensis* monocultures with no supplemental electron source. Encapsulated co-cultures were able to be recycled for more than 30 days with no measurable decrease in graphene oxide reduction efficiency, showing compatibility with future industrial scaling. Photocatalytic hydrogen generation with cadmium selenide quantum dots as the catalyst resulted in 6-fold increases in hydrogen produced by co-cultures using milk as an electron source precursor for the system in comparison to *S. oneidensis* monocultures without any added electron sources. Thus, dairy waste may be processed to drive the synthesis of valuable products utilizing microbial electron transfer processes, converting a significant fluvial environmental pollutant into a valuable renewable energy resource that could provide a robust alternative revenue stream for dairy farmers in a volatile industry.

## Introduction

Biofuels from waste agricultural biomass are a promising option to provide precursor materials for bioprocess reactions, since they can offer decreased reliance on fossil fuels and thus reduce the far-reaching impacts of climate change^6,7^. Previous efforts to develop biofuels have focused on devoting croplands to biofuel production^6^, which can result in environmental degradation and may cause direct and indirect impacts from land use change^6^. Use of food waste that is already generated by the agriculture industry for the production of biofuels could be one route to address these challenges^8^. Cheese whey, a major byproduct of cheese production, accounts for 1.8-1.9 x 10^8^ tonnes of waste per year worldwide^1^. Whey is regarded as environmentally difficult to dispose of due to its high organic content^1^. Utilizing cheese whey for manufacturing high-value products thus creates an opportunity to reduce environmental strain and simultaneously provide extra revenue to dairy farmers. This research presents an opportunity to valorize cheese whey waste by converting it to an electron source to enable microbial production of valuable chemical products, including reduced graphene oxide and hydrogen fuel, using sustainable microbial biotechnologies.

Reduced graphene oxide is a highly conductive material used in thin films for electronics and in energy storage^2,3^. Industrial production of reduced graphene oxide is mostly through the Hummers or Mercano processes, which produce toxic emissions such as hydrazine and NO_X_^2^. Microbial reduction of graphene oxide is a more environmentally-friendly alternative to chemical production^9,10^. *Shewanella oneidensis* (*S. oneidensis*) is a bacterium with the ability to extracellularly reduce electron acceptors, making it a good candidate for microbial reduction of graphene oxide^10^. *S. oneidensis*-reduced graphene oxide has been shown to retain excellent electrochemical properties and high stability^10,11^. For growth and extracellular reduction to occur, an electron source is necessary, which is typically lactic acid or fumarate^12^. However, lactic acid has a high production cost^13^ and large environmental impact^6^ due to the energy cost of its synthesis and the price of precursor chemicals^8^. To reduce these impacts, our research establishes a system in which the agricultural pollutant cheese whey can be converted into lactic acid by bacteria^4,14,15^, which may then be further utilized as a nutrient by *S. oneidensis*, which performs graphene oxide reduction. Lactic-acid-producing bacteria are also adaptable for substrate choice and may consume cardboard^13,16^, food waste^17,18^, or brewer’s waste^16,19^, among others. *Klebsiella pneumoniae* (*K. pneumoniae*) was the lactic-acid-producing bacterium selected for this work as it generates lactic acid from glucose present in cheese waste^20^ and can also act as an electron source for redox chemistry^21,22^.

Implementing a second microbial species into the reduced graphene oxide production platform requires co-culturing the two bacteria in the same reaction vessel, which often poses challenges in implementation. Different species of bacteria may work synergistically in solution^23–26^, but often one bacteria strain will outcompete the other^27^. A few studies have previously established synergistic cultures of *K. pneumoniae* and *S. oneidensis*^28^ to utilize the microbial creation (by *K. pneumoniae*) and consumption (by *S. oneidensis*) of lactic acid^24^. However, these systems required two-pot setups to prevent competition between the bacteria species^29^, and the integration of both strains into a single growth chamber has not been demonstrated. In nature, bacteria can form mutually beneficial multi-species^30^ communities known as biofilms^25,31^ that also offer protection to the bacteria against environmental insults and antimicrobial molecules^31,32^. Bacterial biofilms may be reproduced synthetically by trapping bacteria within a hydrogel structure^27,33,34^. By encapsulating bacterial co-cultures within alginate hydrogel spheroids^33,35,36^, the bacterial colonies are protected, and both species can continue to grow without one strain becoming dominant^26,27,35^.

As a further application for the *S. oneidensis* and *K. pneumoniae* co-culture utilization of dairy waste, this research establishes a circular bioeconomy for hydrogen production by a system comprised of microbes and quantum dots. Hydrogen fuel is a developing renewable energy source that produces only water vapor as waste upon combustion or consumption in fuel cells^37,38^. However, current methods of hydrogen production are energy-intensive and require processing fossil fuels, which contributes negatively to carbon emmisions^38,39^. Microbial hydrogen production, where hydrogen is produced via microbial fermentation or in microbial electrolysis cells, is more sustainable but yields limited output fuel compared to the required input energy^40^. Use of a cadmium selenide (CdSe) quantum dot catalyst to drive hydrogen production photocatalytically was demonstrated^5^ using *S. oneidensis*^41^ respiratory electrons to replenish CdSe in catalysis without the application of an external potential^42^. This photocatalytic system for hydrogen evolution requires lactic acid, which is consumed by the *S. oneidensis* as an electron source^43^ (Figure 1). Introducing *K. pneumoniae* into this photocatalytic bio-nano hybrid system thus allows this microbe to process dairy waste into lactic acid which may be consumed by the *S. oneidensis* bacteria^44^. Intriguingly, *K. pneumoniae* has been demonstrated to produce hydrogen directly by from waste glycerol^45^, giving this microbe the potential to contribute to hydrogen production in the photocatalytic quantum dot platform via multiple pathways.

**Figure 1:**
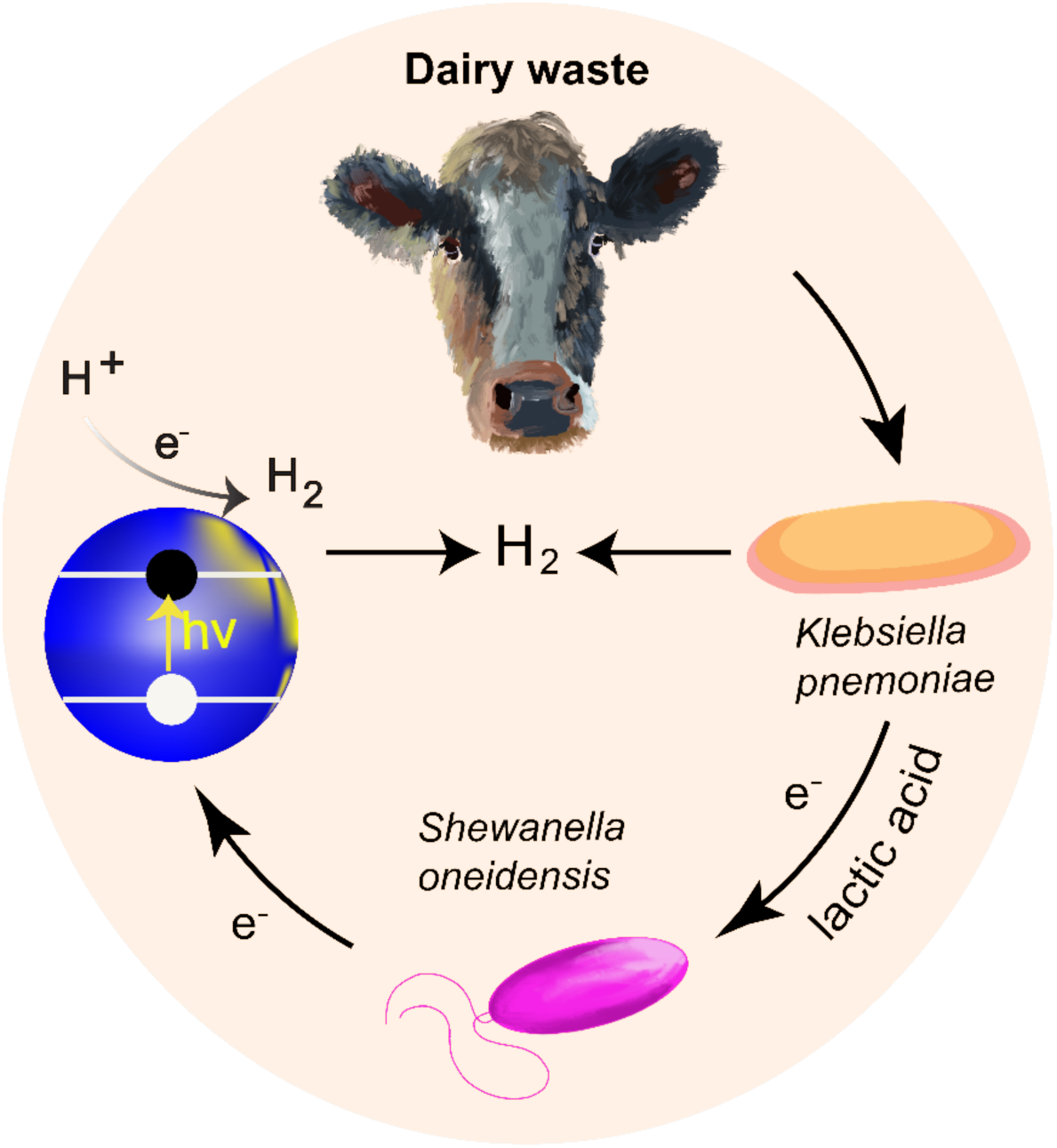
The pathway of hydrogen production for bacterial co-culture systems. *Shewanella oneidensis* consume lactic acid and transfer metabolic electrons to CdSe quantum dot catalysts (blue). Upon illumination at 530 nm, the quantum dots catalyze the production of hydrogen from water. The co-culture utilizes *Klebsiella pneumoniae* to produce lactic acid from glucose. Hydrogen is also produced independently from the hydrogenase pathway of *Klebsiella pneumoniae*.

This work presents a proof-of-concept demonstration of an encapsulated bacterial co-culture system consisting of *K. pneumoniae* and *S. oneidensis*. These bacteria were encapsulated within sodium alginate hydrogel spheroids to generate a one-pot co-culture setup that demonstrated increased bacterial reduction activity and production of reduced graphene oxide and hydrogen fuel utilizing only dairy waste as an electron source precursor. The bacteria-laden alginate spheroids could be re-used multiple times without the need for restarting a new bacterial culture, demonstrating a potential for industrial scalability^33^. The co-culture system could be fueled entirely by unprocessed cheese-making waste, demonstrating a circular economy by converting a significant environmental pollutant into a renewable energy source.

## Results

### Co-cultures of *S. oneidensis* and lactic-acid-producing bacteria can grow using unprocessed dairy waste

To establish a one-pot setup for growing co-cultures of *S. oneidensis* and *K. pneumoniae*, media conditions for the growth of both species were investigated. Wild-type *S. oneidensis* MR-1 cultures, with or without the addition of *K. pneumoniae*, were grown in solution at 30 °C with different media compositions to investigate whether lactose-containing growth media negatively impacted bacterial growth. Bacterial growth was monitored by recording the O.D._600_ of triplicate samples at 10-minute intervals (Figure 2A, B). Maximum growth rates were determined by fitting to a Gompertz growth curve, then determining the slope of a tangent to the inflection point^46–49^ (Figure 2C). Maximum optical density at 600 nm was measured at 20 hours to determine the carrying capacity for growth in solution (Figure 2D). For the growth of *S. oneidensis* in monoculture, cultures with and without lactic acid showed similar growth curves (Fig. 2A), and their maximum growth rates were not significantly different (p=0.42 (0.86,5.6)) (Fig. 2C). The addition of milk or whey resulted in a similar carrying capacity for both (Fig. 2A, D).

**Figure 2:**
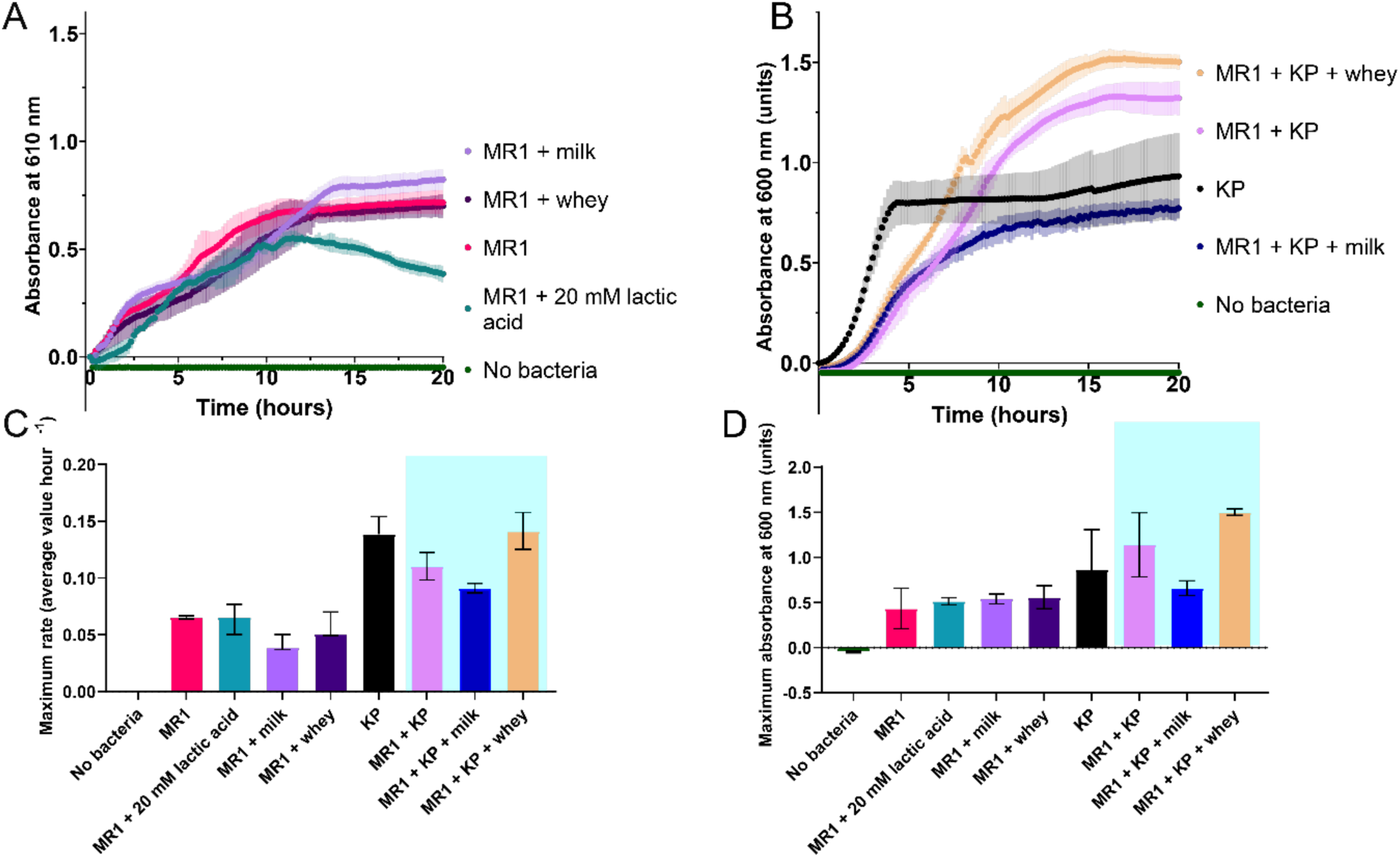
*S. oneidensis* can be co-cultured with *K. pneumoniae* in media containing dairy waste. A, B) The growth of A) *S. oneidensis* (MR-1) monocultures or B) *S. oneidensis*-*K. pneumoniae* (KP) co-cultures in media composed of 1:1 tryptic soy broth (TSB) to lysogeny broth (LB). Media was supplemented with no additions, 20 mM DL-lactic acid, 1% milk powder, or 1% unprocessed whey. *S. oneidensis* were added to media at 1% concentration and *K. pneumoniae* were added at 0.5% concentration from overnight cultures with an initial O.D._600_ of 1. The optical density at 600 nm was monitored at 10-minute intervals in triplicate. No-bacteria samples contained only 1:1 TSB:LB media. C) Maximum growth rates calculated from Gompertz growth fits of the growth curves in A) and B), where the maximum growth rate was estimated by calculating the tangent of the inflection point. D) The maximum values of the growth curves in A) and B) obtained by Gompertz growth fits. All errors represent the standard deviation of triplicate samples.

For *K. pneumoniae* and *S. oneidensis* co-cultures, the sum of the maximum growth rates of the two monocultures (0.21 ± 0.02 units hour^−1^) was higher compared to that of the co-culture (0.12 ± 0.02 units hour^−1^) (Fig. 2C), and the summed maximum optical density values of the monocultures (1.7 ± 0.15 units) was higher than that of the co-culture (0.88 ± 0.28 units) (Fig. 2D). These results indicated the occurrence of interspecies competition between strains. The maximum optical density value of the co-culture was not significantly different from the co-culture with milk as an additive (p=0.1065 (1.887,5.177)) but was significantly lower than co-culture growth with whey as an additive (p=0.0019 (5.669,5.354)) (Fig. 2D). This result implied that the whey was being utilized by the co-culture as additional nutrients in addition to the growth media. Monitoring of lactic acid production by *K. pneumoniae* indicated that it produced the most acid during anaerobic growth, followed by aerobic growth in TSB media. Acid production was the lowest during growth in LB or minimal media (Supplemental Figure 1)^50^, although acid production was still detected in these non-ideal growth conditions. Collectively, these results demonstrated that non-ideal growth conditions in the presence of dairy waste can support lactic acid production by *K. pneumoniae* and co-culture establishment, but interspecies interactions can variably impact growth kinetics and biomass yields.

### Enhanced extracellular electron transfer activity of *S. oneidensis* and *K. pneumoniae* co-cultures in solution

To observe whether co-culture growth allowed for enhancement of *S. oneidensis* reduction activity in the presence of milk or whey as an electron source precursor, the reduction activities of co-cultures of *S. oneidensis* and *K. pneumoniae* in solution were compared to monocultures. An assay observing the reduction of graphene oxide^51^ over time was used for initial testing of microbial reduction activity under various co-culture conditions^11,52^ ^42^.

Cultures were supplemented with graphene oxide, and optical density at 610 nm was measured over time to detect the appearance of reduced graphene oxide as GO displays an emission maximum at this wavelength in media of neutral pH^53^(Fig. 3A). Since the optical density of the bacteria within the samples partially overlapped with the optical density of the reduced graphene oxide at 610 nm^52^, the average optical density of bacteria samples that did not contain graphene oxide was subtracted from each corresponding sample. Graphene oxide addition has been observed to result in minimal negative impact on *S. oneidensis* growth^54^ but could contribute to a slight underestimation of reduction.

**Figure 3:**
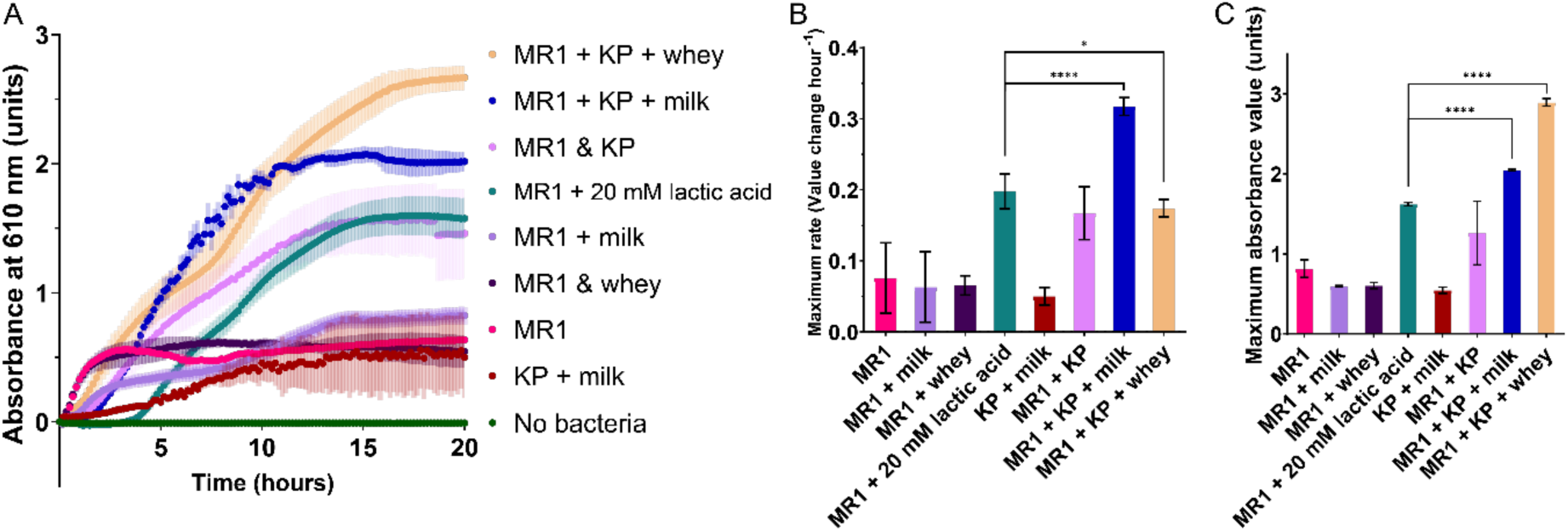
Solution-based co-cultures show enhanced graphene oxide reduction activity compared to monocultures. A) Reduction of graphene oxide over time was monitored by optical density at 610 nm for co-cultures of *S. oneidensis* (MR-1) with *K. pneumoniae* (KP) in a 2:1 ratio. Co-cultures were grown in 1:1 TSB:LB media with graphene oxide, with 1% milk powder, 1% unprocessed whey, or 20 mM DL-lactic acid added to solution as indicated. O.D._610_ measurements of the same samples not containing graphene oxide have been subtracted from each data set to correct for overlap from bacterial optical density. B) The maximum reduction rates calculated from Gompertz growth fits of the reduction reactions for the co-cultures, where the maximum reduction rate was estimated by calculating the tangent of the inflection point. C) The maximum optical density value obtained by Gompertz growth fits, indicating the total amount of reduction. All error bars represent the standard deviation of triplicate samples. * P≤0.05, **** P≤0.0001 by one-way ANOVA statistical analysis and comparative two-tailed t-tests performed with a Welch’s correction.

The monocultures containing only *S. oneidensis* with no supplemented electron source displayed a modest rate of reduction which was likely due to usage of components in the nutrient medium as poor-quality electron sources^55–57^ (Fig. 3B). The change in O.D._610_ over time, approximated as representing the reduction rate of these monocultures of *S. oneidensis* with no supplemented electron source, was not significantly different from the reduction rate of *S. oneidensis* monocultures with added milk (p = 0.42 (0.93,3.1)) or added whey (p = 0.28 (1.2, 4.8)) (Fig. 3B). The total reduction of graphene oxide by the monocultures of *S. oneidensis* with no supplemented electron source was not significantly different from the total graphene oxide reduction by *S. oneidensis* monocultures with added milk (p = 0.072 (3.46,2.03)) or whey (p = 0.071 (3.24,2.43)) (Fig. 3C). These data indicate that the *S. oneidensis* was not significantly utilizing the unprocessed milk or whey itself as an electron source for graphene oxide reduction. The total graphene oxide reduction of the *S. oneidensis* monoculture with added lactic acid as an electron source (Fig. 3C) was significantly different from *S. oneidensis* monoculture with added whey (p < 0.0001 (51.3, 6.72)) or added milk (p < 0.0001 (76.4, 2.85), indicating that supplemented electron source is required for efficient graphene oxide reduction and that the whey and milk in their unprocessed forms are insufficient as an electron source for the *S. oneidensis*. Interestingly, the co-culture with no added milk or whey had a graphene oxide reduction rate comparable to the monoculture of *S. oneidensis* with lactic acid (p = 0.506 (2.93,3.49)) and a comparable total reduction (p=0.257 (1.57,2.01)), indicating that the *K. pneumoniae* was likely processing a component of the nutrient medium into a form that could be utilized for reduction. The *K. pneumonaie* monoculture with milk as an additive did perform measurable reduction, but the reduction rate was small and not significantly different from that of the monoculture of *S. oneidensis* with no added electron source (p=0.15 (2.1,2.2)).

Co-cultures of *S. oneidensis* and *K. pneumoniae* with added milk or whey had significantly increased maximum rates of graphene oxide reduction compared to the *S. oneidensis* monoculture with no additives (Fig. 3B). Compared to co-cultures in unsupplemented media, co-cultures with added milk showed a 420% increase (p = 0.0014 (20.30,2.249)), and those with added whey showed a 230% increase (p= 0.010 (8.27,2.24)) in maximum reduction rate. Similarly, total reduction was also increased for co-cultures, with added milk resulting in a 250% increase (p = 0.003 (19.3,2.03)) and added whey causing a 354% increase (p < 0.001 (30.1,2.71)). These results are consistent with processing of the milk or whey by the *K. pneumoniae* into lactic acid to provide a supplemental electron source for the *S. oneidensis* to drive extracellular reduction. Co-cultures of *S. oneidensis* and *K. pneumoniae* showed significantly increased maximum rates of reduction compared to the MR-1 monoculture with lactic acid added, when milk (160% increase, p = 0.0004 (18.50,2.91)) or whey (13% increase, p = 0.0361 (3.675, 2.941)) was added. Similarly, the total reduction was also increased for the co-cultures with milk (26% increase, p <0.0001 (31.87,2.863)) and whey (79% increase, p < 0.0001 (42.87, 2.768)) as an additive. These increases may be attributable to robust, continual production of lactic acid by the co-cultured *K. pneumoniae* compared to the limited lactic acid that was supplied in the MR-1 culture medium. These results demonstrated a significant improvement in the reduction activity of this co-culture when fed with dairy products, despite negative growth impacts from competition of the species within the solution (Fig. 2). The enhancement in reduction seen for the co-cultures compared to the low reduction activity exhibited by both individual monocultures in the presence of milk was likely due to production of lactic acid by the *K. pneumoniae* that could serve as a supplemental electron source to fuel the reduction activity of the *S. oneidensis*.

### Enhanced reduction activity of alginate-encapsulated co-cultures

It was hypothesized that reducing competition between bacterial species through physical separation via alginate encapsulation would increase the graphene oxide reduction activity of the co-cultures. To test this hypothesis, individual bacteria cultures were resuspended in alginate-based bio-ink^58–61^ and extruded with a syringe pump overtop of a calcium chloride bath to produce alginate spheroids containing encapsulated bacteria cells (Supplemental Figure 2). The spheroids displayed consistent diameters, with an average value of 0.422 ± 0.086 cm (Supplemental Figure 3). The narrow spread of measurements demonstrated that the spheroid fabrication process produced a homogenous population suitable for downstream co-culture assays.

Spheroids containing different bacteria species were individually added to the same growth media to allow the bacteria species to be co-cultured in the same culture vessel (Fig. 4A). To determine the effect of alginate spherification on the reduction activities of the co-cultures, graphene oxide reduction by the spherified bacteria was measured. Images of the samples (Supplemental Figure 4) were processed using a recently developed image analysis method^52^ to calculate the average HSV value difference after 23 hours (Fig. 4B) as a measure of graphene oxide reduction. The negative values measured for the samples containing no bacteria reflected a decrease in optical density of the graphene oxide over time due to dispersal within the media^3,62^. Some reduction was detected for the *S. oneidensis* utilizing milk as an added electron source, but this reduction activity was significantly lower compared to the *S. oneidensis* monoculture with lactic acid as an electron source (p = 0.0089 (3.192,10.70)) (Fig. 4B). The monoculture of *K. pneumoniae* with milk as an additive also produced measurable graphene oxide reduction that was not significantly different from the *S. oneidensis* monoculture with lactic acid as an electron source (p = 0.1224 (1.626,17.00)) (Fig. 4B). When compared to the *S. oneidensis* monoculture, the *K. pneumoniae* and *S. oneidensis* co-culture showed significantly higher reduction activity with the addition of milk (333% increase, p <0.0001 (6.439, 19.98)) or whey (421% increase, p < 0.001 (7.995,19.48)) (Fig. 4B), indicating processing of the milk or whey by the *K. pneumoniae* into an electron source, likely lactic acid, for the *S. oneidensis.* The *K. pneumoniae* and *S. oneidensis* co-culture with milk as an additive showed significantly higher reduction activity than the *S. oneidensis* monoculture with lactic acid as an electron source (p = 0.0083 (2.927, 19.99)), as did the co-culture with whey as an additive (p = 0.0001 (4.867,19.13)) (Fig. 4B). When co-cultures were combined within the same spheroids rather than separated, no significant difference in cumulative reduction was measured between the two groups (Supplemental Fig. 5), demonstrating that direct mixing of cultures within a spheroid did not enhance extracellular substrate reduction compared to spatially separated co-cultures. These results indicate successful processing of lactose by the *K. pneumonaie* into lactic acid for utilization as an electron source by the *S. oneidensis*.

**Figure 4:**
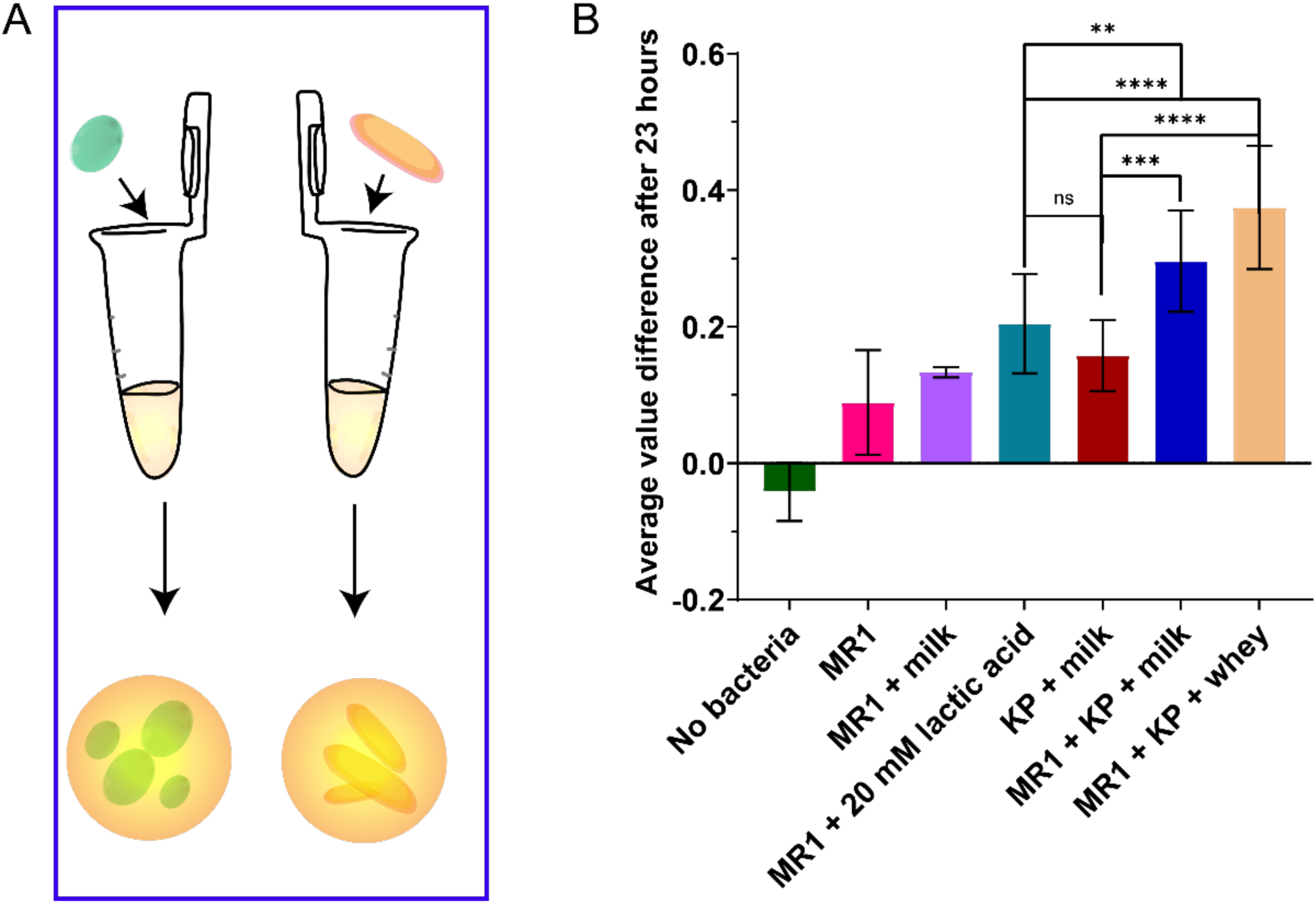
Encapsulated bacterial co-cultures demonstrate improved extracellular reduction relative to monocultures. A) *S. oneidensis* and *K. pneumoniae* cultures were encapsulated in separate alginate spheroids. B) Spheroids containing monocultures of *S. oneidensis* and *K. pneumoniae* were added to minimal media containing graphene oxide with added 20 mM lactic acid, 1% milk powder, or 1% whey. Images were taken at 0 hr and 23 hr timepoints, and the average change in HSV value of each sample was calculated Error bars represent the standard deviation of triplicate samples. ** p≤0.01, *** p≤0.001, **** p≤0.0001, ns: not significant by one-way ANOVA statistical analysis and two tailed t-tests performed with a Welch’s correction.

### Long-term reusability of alginate-encapsulated co-cultures with enhanced reduction ability

To establish the applicability of the alginate-encapsulated bacterial spheroids (Fig. 4A) for future bioreactor compatibility the reusability of the spheroids was examined over a period of 32 days. Spheroids containing co-culture species were placed into fresh media containing graphene oxide, and the cultures were imaged at time 0 and 23 hours to obtain the difference in HSV value. The spheroids were removed and placed into new media thrice a week, whereupon the samples were then imaged at 0 and 23 hour timepoints again. The cumulative amount of HSV value difference reflects the overall amount of extracellular reduction over time for this 32-day experiment (Fig. 5A). When spheroids contained mixtures of both co-culture species as opposed to the species being separated within different spheroids, the differences in reduction between the mixed and separated samples were non-significant over the time period of the experiment (Supplemental Figure 6), indicating that the overall reduction capacity in this set-up was primarily due to the protective alginate encapsulation rather than the relative arrangement of the two bacteria species in the spheroids.

**Figure 5:**
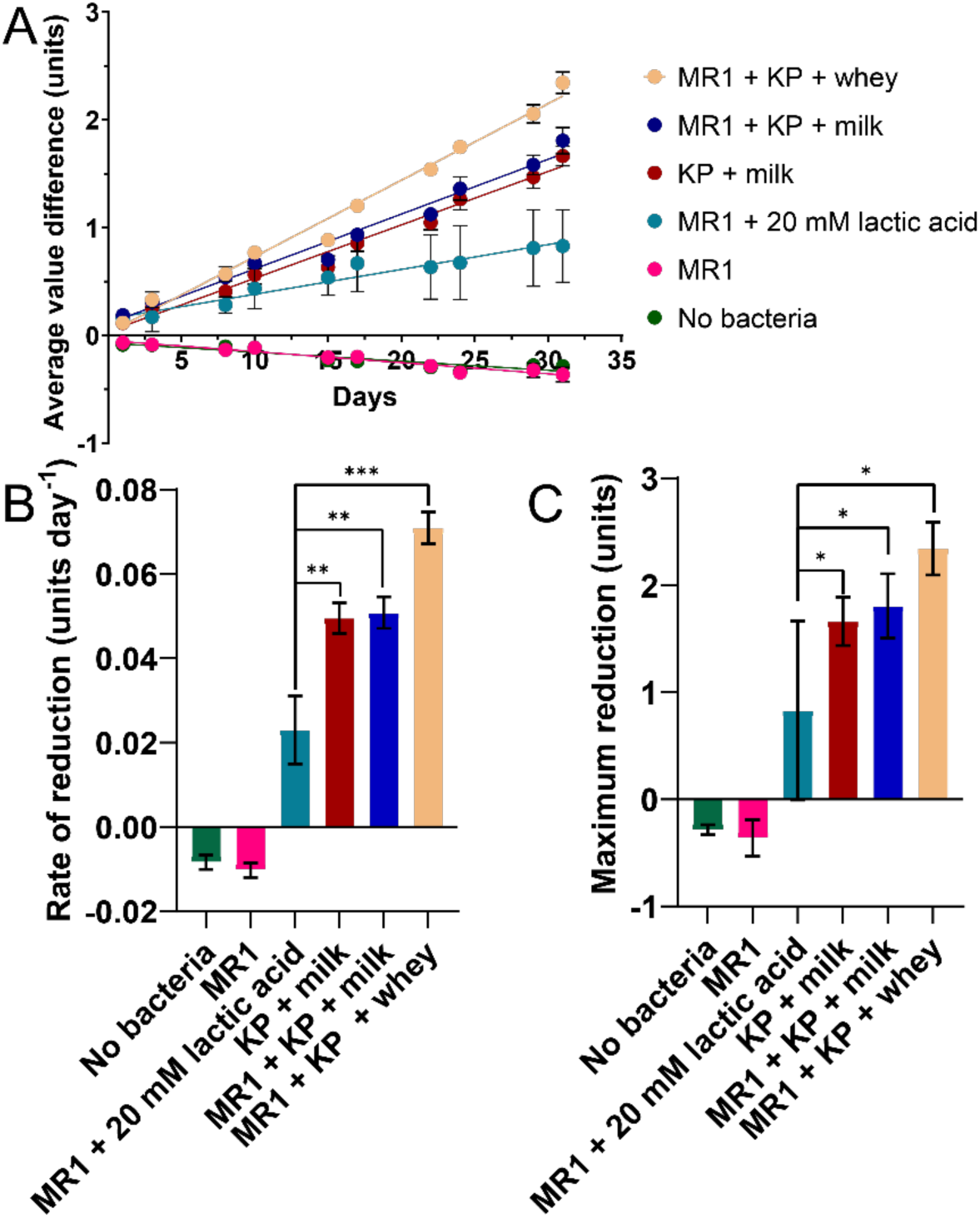
Extended re-usability of encapsulated co-cultures. A) *S. oneidensis* and *K. pneumoniae* cultures were encapsulated in separate alginate spheroids and added to minimal media containing graphene oxide with added 20 mM lactic acid, 1% milk powder, or 1% whey. Images were taken at 0 hr and 23 hr timepoints, and the average HSV value of each sample was calculated. Spheroids were transferred into fresh media for every subsequent measurement, three times a week over 32 days. The cumulative average value differences over time are presented. B) The average rate of reduction and C) the maximum cumulative reduction over the course of the 32 days. Error bars represent the standard deviation of triplicate samples. * p≤0.05, ** p≤0.01, *** p≤0.001 by one-way ANOVA statistical analysis and two-tailed t-tests performed with a Welch’s correction.

For experiments using spheroids that contained individual single bacteria species, all monocultures and co-cultures tested showed a consistent, steady rate of graphene oxide reduction throughout the 32-day time course, as demonstrated by the linear increase of reduction over time (Fig. 5A). The spherified *K. pneumoniae* and *S. oneidensis* co-cultures showed enhanced reduction rates compared to the *S. oneidensis* monocultures both with and without lactic acid as an added electron source (Fig. 5B). When compared to the *S. oneidensis* monoculture with no added electron source, a significant increase was observed in the average rate of reduction of the milk-fed (p <0.0001 (52.51,2.783)) and the whey-fed co-cultures (p < 0.0001 (69.85, 2.785)) (Fig. 5B). The total amount of reduction was similarly increased for the milk-fed (p < 0.0001 (27.00,3.152)) and whey-fed (p < 0.0001 (38.82, 3.450)) co-cultures (Fig. 5C). The rate and total amount of the reduction of the *S. oneidensis* monoculture with no supplemented electron source was negative since graphene oxide will disperse in media over time^52^, decreasing the HSV value recorded by imaging. The large increase in reduction measured for the co-cultures with added milk or whey compared to the *S. oneidensis* monoculture demonstrated that successful processing of the lactose by the *K. pneumonaie* was maintained over an extended period of time. The milk-fed co-cultures displayed a 120 % increase in the reduction rate (p=0.0020 (11.09,2.832)), and the whey-fed co-cultures showed a 208 % increase (p=0.0004 (19.11,2.830)) compared to the *S. oneidensis* monoculture supplemented with lactic acid (Fig. 5B). Interestingly, the *K. pneumoniae* monoculture fed on milk also displayed a significantly higher reduction rate (0.050 ± 0.003 units day^−1^) than the *S. oneidensis* monoculture supplemented with lactic acid (0.023 ± 0.008 units day^−1^) (p=0.0025 (10.63,2.774)) (Fig. 5B). Maximum reduction was significantly increased for the milk-fed co-culture (p=0.0263 (4.737,2.511)) and the whey-fed co-culture (p=0.0109 (7.479,2.348)) when compared to the *S. oneidensis* monoculture with or without supplemented lactic acid (Fig. 5C). These data indicated that alginate-encapsulation of co-culture strains allowed for re-usable, enhanced extracellular electron transfer by *S. oneidensis* and *K. pneumoniae* co-cultures using milk or dairy waste as an additive.

### Photocatalytic hydrogen production by spherically encapsulated co-cultures with CdSe quantum dots utilizing dairy waste

To observe the impact of co-cultures on hydrogen production in a system with colloidal CdSe quantum dots (QDs), the alginate-encapsulated bacteria were incubated with CdSe QDs and evolved hydrogen was measured. CdSe QDs were coated with cysteine ligands to ensure their stability and solubility in water^63^. Alginate-encapsulated bacterial strains were added in a 2:1 ratio of *ΔhydAΔhyaB S. oneidensis* to *K. pneumoniae* and were compared against monocultures and control conditions (Fig. 6A). In place of wild-type *S. oneidensis* MR-1, a hydrogenase knockout *ΔhydAΔhyaB S. oneidensis* strain was used to prevent both direct microbial production of hydrogen through non-extracellular electron transfer pathways as well as the metabolism of hydrogen by *S. oneidensis* at high concentrations^64^.

**Figure 6:**
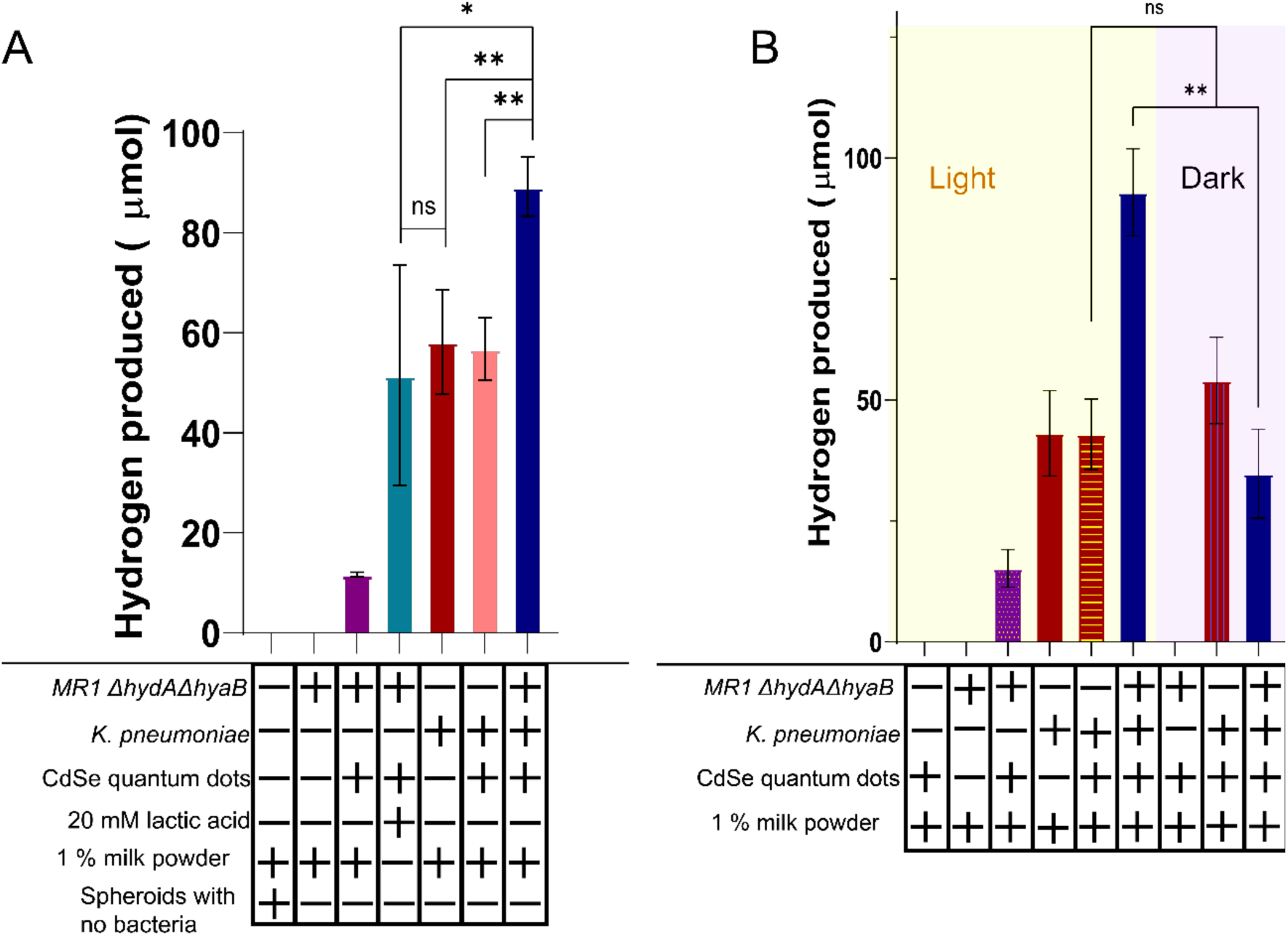
Hydrogen production is increased for spherified co-cultures. A) Hydrogen production by co-cultures and monocultures of spherified *ΔhyaAΔhydB S. oneidensis* and *K. pneumoniae* was tested with and without 1% milk powder, CdSe quantum dots, 20 mM lactic acid, light illumination, and alginate spheroids containing no bacteria. B) Hydrogen production of the spherified co-cultures and monocultures both with (Light) and without (Dark) illumination. Error bars represent the standard deviation of triplicate samples. * p≤0.05, ** p≤0.01, ns: not significant by one-way ANOVA statistical analysis and two-tailed t-tests performed with a Welch’s correction.

No hydrogen production was detected when *ΔhydAΔhyaB S. oneidensis* monocultures were incubated with milk and light illumination and no QDs, indicating successful inactivation of the hydrogenase enzymes in the mutant strain. A small amount of hydrogen was produced from illuminated samples containing *ΔhydAΔhyaB S. oneidensis*, milk, and QDs, consistent with a minor contribution to hydrogen production from *S. oneidensis* utilizing unprocessed milk as an additive. Since the illuminated sample containing QDs and 1% milk with no microbes produced no measurable hydrogen, it is unlikely that the milk was acting directly as an electron source for the CdSe QDs. The illuminated *K. pneumoniae* monocultures produced similar, high amounts of hydrogen both in the presence (43 ± 7 µM hydrogen) and absence of CdSe QDs (43 ± 9 µM hydrogen) (Fig. 6A), suggesting that this production occurs along a microbial hydrogenase pathway^65,66^ rather than from catalytic activity of the QDs. The *K. pneumoniae* also produced similar levels of hydrogen when there was no light illumination (54 ± 8.9 µM hydrogen) (Fig. 6B), further supporting this hypothesis.

Illuminated co-culture solutions containing CdSe QDs with milk as an additive produced significantly more hydrogen than *S. oneidensis* monocultures with CdSe QDs and lactic acid (p=0.046 (2.86,4.00)) as an electron source, showing a 173% increase in produced hydrogen (Fig. 6A). In comparison to the illuminated *S. oneidensis* monocultures with CdSe QDs and milk, the co-cultures incubated with QDs and milk showed a 665% increase in hydrogen production (p = 0.002 (22.4,2.02)) (Fig. 6A). This result implies that *K. pneumoniae* was able to convert the milk into components that were utilized by the *S. oneidensis* bacteria to support hydrogen production by the catalytic nanoparticles. The summed total of the hydrogen production of the illuminated *S. oneidensis* monocultures incubated with QDs and milk and the illuminated *K. pneumoniae* monocultures incubated with QDs and milk was 67 ± 6 µM hydrogen, whilst the total hydrogen produced by the co-cultures incubated with quantum dots with milk as an additive was 89 ± 6 µM hydrogen. This synergistic increase is consistent with the *S. oneidensis* utilizing lactose processed into lactic acid by the *K. pneumoniae* as an electron source to increase the extracellular reduction of hydrogen.

The co-culture reaction with milk as an additive produced significantly higher levels of hydrogen under light illumination (93 ± 9 µM hydrogen) than without (35 ± 9 µM hydrogen) (p=0.0014 (7.835,3.999)) (Fig. 6B). This result indicated that the production of hydrogen was dependent on the extracellular transfer of electrons by *S. oneidensis* to the catalytic CdSe QDs, similar to previous results shown for *S. oneidensis* monocultures^64^. The hydrogen produced by the co-culture, milk, and CdSe QD system without illumination was not significantly different from the level of hydrogen produced microbially by the *K. pneumoniae* without light illumination (p=0.1723 (1.674, 3.845, Fig. 6B), indicating that the residual hydrogen was likely produced by the *K. pneumoniae* through its hydrogenase pathway. Thus, *K. pneumoniae* not only generated hydrogen through direct metabolism but also by processing the milk to enable further hydrogen production by the *ΔhydAΔhyaB S. oneidensis.* Overall, the highest amount of hydrogen production was seen for solutions containing both *ΔhydAΔhyaB S. oneidensis* and *K. pneumoniae* incubated with QDs and milk in the presence of light. The co-cultures allowed enhancement of hydrogen production compared to either monoculture, likely due to a combination of direct hydrogen production by the *K. pneumoniae* and the production of lactic acid from milk by the *K. pneumoniae* as an electron source to support quantum dot reduction by the *S. oneidensis*.

## Discussion

This study demonstrates that alginate-encapsulated co-cultures of *S. oneidensis* with *K. pneumoniae* are an effective technology for increasing extracellular electron transfer for reduced graphene oxide production. We demonstrate that whey, a type of unprocessed cheese waste, can be utilized as an electron source after microbial processing to lactic acid.

Collectively, our results support the potential of the *S. oneidensis* and *K. pneumoniae* co-culture to contribute to a circular bioeconomy approach for dairy waste utilization for both graphene oxide reduction and hydrogen fuel production. In solution, interspecies competition was seen to have negative impacts on co-culture performance. However, encapsulation within alginate matrices enhanced co-culture performance, likely by buffering detrimental environmental fluctuations^67^ and limiting competitive pressures. Encapsulated cultures maintained stable reduction activity over 32 days, indicating compatibility with continuous industrial operation. These results establish a scalable framework for integrating waste-to-fuel conversion with nanomaterial production and offer a new direction for sustainable industrial bioprocess development.

Towards the creation of a circular bioeconomy, we observed whether the lactic acid produced by the *K. pneumoniae* was utilized by the *S. oneidensis* for extracellular reduction, using the reduction of graphene oxide as a model assay^53,63^. Reduced graphene oxide is valued for electronic and optical applications due to its biocompatibility and conductivity^68,69^, while its chemical synthesis poses environmental and economic concerns, making efficient microbial reduction highly desirable^68^. The graphene oxide reduction rate for the *K. pneumoniae* and *S. oneidensis* co-culture was significantly improved from the monocultures despite expected interspecies competition, showing a marked increase in maximum reduction amount for both the milk-fed (2.5 fold increase in total reduction) and the whey-fed co-culture (3.5 fold increase in overall reduction) when compared to the *S. oneidensis* monoculture with no supplemental electron source. A small amount of graphene oxide reduction was performed by the monoculture of *K. pneumoniae* with milk as an additive, although this result was not significantly different from the reduction performed by the *S. oneidensis* monoculture with no additive electron source. Some chemical reduction caused by *K. pneumoniae* electron shuttling has been previously reported, although the direct pathway has not been fully investigated^44,70^. Collectively, these findings indicated successful processing of the milk or whey by the *K. pneumoniae* into a form, likely lactic acid, that could be utilized by the *S. oneidensis* as an additive electron source for extracellular reduction of the graphene oxide.

To further improve the graphene oxide reduction of the co-culture, it was hypothesised that physical separation by alginate encapsulation of the individual microbe species may alleviate potential pH^67,71,72^ and toxin^72,73^ effects from localized lactic acid production by the microbes as well as reducing interspecies competition^74–77^. Alginate encapsulation would also enable nutrient media to be flowed through a bioreactor without having to restart the microbial culture^72^, which could enable future industrial scaling^78^. Notably, the alginate-encapsulated spheroids demonstrated long-term reusability, with consistent and linear reduction activity maintained for over 30 days (Figure 5), highlighting their promise for industrial graphene oxide manufacturing applications^72^. In contrast to the solution-based cultures, greater enhancements in total reduction were observed with *K. pneumoniae* and *S. oneidensis* co-cultures, which displayed 333% and 421% increases under milk- and whey-fed conditions, respectively compared to *S. oneidensis* monocultures with no supplemental electron sources. Interestingly, *K. pneumoniae* monocultures grown on milk outperformed *S. oneidensis* monocultures provided with lactic acid as an added electron source with regards to both graphene oxide reduction rate and total amount of reduction, suggesting a greater-than-anticipated extracellular reduction capacity for *K. pneumoniae*^44,70^.

The improved performance of the *K. pneumoniae* and *S. oneidensis* co-culture with regards to increased graphene oxide reduction rate and overall reduction may reflect synergistic interactions, wherein *K. pneumoniae* supplies lactic acid as an electron donor for *S. oneidensis* ^79^ while simultaneously reducing graphene oxide directly. The strong reduction activity of whey-fed co-cultures was unexpected, as *K. pneumoniae* predominantly metabolizes glucose^79,80^. Lactose fermentation by *K. pneumoniae* has been previously reported^81,82^, though its metabolic byproducts have not been characterized in detail. Prior studies on *K. pneumoniae* biofuel production have largely focused on metabolism of glucose^66,79^ or glycerol^83,84^, and to our knowledge, utilization of unaltered cheese waste has not been previously examined. These findings suggest that further investigation into the metabolism of whey by *K. pneumoniae* could reveal additional opportunities to optimize biofuel production from dairy sources. The enhanced reduction rates achieved with alginate-encapsulated co-cultures highlight the importance of the alginate to decrease interspecies competition, consistent with previous findings using 3D-printed co-cultures^78^. Overall, these results support the feasibility of scaling up alginate-encapsulated microbial systems for reduced graphene oxide generation directly from unprocessed cheese whey, sustainably converting an environmental pollutant into a high-value product.

As an additional application, the ability of the alginate-encapsulated *K. pneumoniae* to process milk into lactic acid was leveraged to fuel *S. oneidensis* extracellular transfer of electrons to CdSe quantum dots, producing hydrogen. Remarkably, *K. pneumoniae* monocultures exhibited equivalent hydrogen production to *S. oneidensis* monocultures supplemented with lactic acid as an electron source, suggesting substantial direct hydrogen production by the *K. pneumoniae*. There has been sparse investigation into *K. pneumoniae* extracellular reduction activity, but an electron shuttle mechanism^44^ and a NADH dependent pathway^65,85^ have been reported^64,83,86^. The hydrogen production was not significantly different between *K. pneumoniae* monocultures with or without light illumination, indicating that the extracellular electron transfer was quantum-dot-independent and thus likely occurs through a hydrogenase pathway within the *K. pneumoniae*^80,83^. Co-cultures of *K. pneumoniae* and *S. oneidensis* containing QDs and milk as an additive produced a 6.6-fold increase in hydrogen when compared to *S. oneidensis* monocultures with quantum dots and milk as an additive. This result indicated *S. oneidensis* utilization of the lactic acid microbially generated by the *K. pneumoniae* from the added milk. The co-culture hydrogen production was higher than the additive sum of the *K. pneumoniae* monoculture with milk and the *S. oneidensis* monoculture with no supplemented electron source. This result implies that the co-culture successfully utilized the lactose processed by the *K. pneumoniae*, but that the increased hydrogen production from the co-culture was in part due to the hydrogen produced independently by the *K. pneumoniae.* These results demonstrate that *K. pneumoniae* not only supplied fermentative metabolites to support *S. oneidensis* respiration, but also directly produced hydrogen, enabling effective conversion of complex dairy waste streams into hydrogen fuel. Together, these findings establish a novel platform that can valorize cheese waste into high-value products such as reduced graphene oxide and hydrogen fuel. This co-culture system shows compatibility with future industrial scalability, advancing prospects for waste valorization on a large scale.

## Experimental methods

### Growth of Shewanella oneidensis

*S. oneidensis MR-1* (ATCC® 700550™) cultures or *S. oneidensis ΔhydAΔhyaB* hydrogenase knockout cultures (kindly provided by Dr. Benjamin Keith Keitz and Dr. Jeffrey Gralnick^86^) were grown from freezer stocks stored in 25% glycerol at −80 °C. Lysogeny broth (LB) media was prepared by dissolving tryptone (0.14 M, Sigma-Aldrich), yeast extract (15 mM, Sigma-Aldrich), and NaCl (0.17 M, Sigma-Aldrich) in deionized water. This solution was autoclaved for 30 minutes on a fluid cycle at 121 °C and stored at room temperature. Frozen bacterial cultures were streaked onto LB-agar plates and incubated at 30 °C overnight. The following day, a single colony was isolated and grown in 5 mL LB at 30 °C overnight under continuous shaking (200 rpm). These overnight cultures were used for further experiments.

### Growth of Klebsiella pneumoniae

*K. pneumoniae* (Ward’s live *Klebsiella pneumoniae* culture, Rochester, NY, USA) cultures were grown from freezer stocks stored in 25% glycerol at −80 °C. Tryptic soy broth (TSB) media was prepared by dissolving tryptic soy broth powder (20 g L^−1^, Sigma-Aldrich) in deionized water. This solution was autoclaved for 30 minutes on a fluid cycle at 121 °C and stored at room temperature. Frozen bacterial cultures were streaked onto LB-agar plates and incubated at 37 °C overnight. The following day, a single colony was isolated and grown in 5 mL TSB at 37 °C overnight under continuous shaking (200 rpm). These overnight cultures were used for further experiments.

### Preparation of milk and whey

Skimmed milk powder (10 g L^−1^, Judee’s, USA) was dissolved in deionized water. This solution was heated to 65 °C for one hour to sterilize the milk and avoid bacterial contamination. The skimmed milk solution was streaked onto LB-agar plates and incubated overnight at 30 °C to confirm a lack of bacterial contamination. Waste whey product from goat cheese making was obtained from Pete Messmer at Lively Run Goat Dairy (Interlaken, NY) in an unaltered state and stored at −80 °C. To avoid contamination by bacteria remaining from the cheese-making process, the whey was heated to 100 °C for 1 hour and then cooled to room temperature before use.

### Color analysis of graphene oxide reduction sample images

Images of graphene oxide reduction were analyzed using the technique published previously in Bennett and Meyer (2025)^87^ with minor addendums. Briefly, each image was taken using a Google Pixel^TM^ phone using a RAW image format to avoid contrast adjustment by the Google software. Samples were illuminated from below using a miroco SAD lamp (miroco) for even illumination. For each image, the sample was selected and cropped to select an area that had minimal interference from background or reflections. This area was converted to HSV format, which represents the color information of the selected area using three vectors^88^. The HSV color space is advantageous over other color spaces for image analysis, such as the more typical RGB, because the data obtained is more stable under changing lighting conditions^89^. The V vector contains value information: how light or dark each pixel is. The value vector was isolated, and the average value was taken over the entire box. Each pixel was represented by a number between 0 and 1, where 0 is the darkest and 1 is the lightest.

### Growth curves and lactic acid measurement

A single bacteria colony was selected and grown overnight in LB media (*S. oneidensis*, 30 °C) or TSB media (*K. pneumoniae*, 37 °C). The O.D._600_ was measured for each overnight culture, and cultures were adjusted to a concentration of 1 O.D._600_. This culture was added to a 24-well tissue culture plate (Fisherbrand, Thermo-Fisher Scientific) containing the growth medium to be examined (LB or TSB media), to a final concentration of 1%. Milk solution was added to 1% concentration (0.1 g L^−1^). Anaerobic samples were prepared by sealing the solution in an EPA vial and flushing with nitrogen gas for 10 minutes. After transfer to the 24 well tissue culture plates, the wells were covered with mineral oil (500 µL, Sigma-Aldrich) to maintain seclusion from oxygen. Wells were monitored at 10-minute time intervals at a wavelength of 600 nm in a Biotek© Synergy H1 microplate reader to observe the bacterial growth. Aliquots (0.5 mL) were removed from the total solution volume (20 mL) at timepoints to monitor the pH of the solution using a VWR pH probe.

### Reduction activity of *K. pneumoniae* and *S. oneidensis* in solution

A single bacterial colony was selected and grown overnight in LB media (*S. oneidensis*) or TSB media (*K. pneumoniae*). The O.D._600_ was measured for each overnight growth media and was adjusted to a concentration of 1 O.D._600_. This culture was added to a 24 well tissue culture plate (Fisherbrand, Thermo-Fisher Scientific) containing the growth medium (1:1 ratio of LB:TSB media)) for a final concentration of 1%. Graphene oxide (0.05 g L^−1^, Sigma-Aldrich) was added to each well. Anaerobic samples were prepared by sealing the solution in an EPA vial and flushing with nitrogen gas for 10 minutes. After transferral to the 24 well tissue culture plates, the wells were covered with mineral oil (500 µL, Sigma-Aldrich) to maintain seclusion from oxygen. Wells were monitored at 10-minute time intervals at a wavelength of 610 nm in a Biotek© Synergy H1 microplate reader to observe the bacterial growth at a temperature of 35 °C.

### Preparation of bacteria and alginate bio-ink

Overnight bacteria cultures were centrifuged at 3000 xg for 15 mins to pellet the bacteria. Sodium alginate solution was prepared by dissolving sodium alginate (3 g L^−1^) in a solution of 1:1 (LB:TSB) medium and heating to 100 °C while stirring. The bacterial pellets were resuspended in sodium alginate solution for a final O.D._600_ of 20.

### Preparation of bacterial alginate spheroids

Bacteria and alginate bio-ink were prepared as outlined above. Calcium chloride (0.1 M) was added to LB media. The bacterial alginate solution was collected using a 10 mL syringe (Exel 26265). A syringe pump (New Era Pump Solutions Inc.) was used to drop the bacterial alginate solution into the calcium chloride solution at a speed of 5.10 mL / hr, forming spheroids of uniform diameter (SI Figure 4). The spheroids were left in solution for 10 mins to allow full crosslinking of the alginate. Spheroids were then transferred to a solution of 1:1 LB:TSB with calcium chloride (0.05 M) and incubated at 30 °C overnight before use. For longer term storage, spheroids were stored at 30 °C, and the calcium chloride solution was refreshed every two days.

### Reduction by re-used co-cultures encapsulated in alginate spheroids

Spheroids were prepared as described above containing *S. oneidensis, K. pneumoniae,* or combinations of *S. oneidensis* and *K. pneumoniae* bio-inks. Spheroids (1 of each separate bacterial species, or 2 for the experiments with species blended within the spheroids) were added to 1 mL of media (1:1 TSB:LB media) in individual wells in a 24-well tissue culture plate (Fisherbrand, Thermo-Fisher Scientific). Graphene oxide was added to each well to a final concentration of 0.05 g L^−1^. Calcium chloride was added to each well to a final concentration of 0.05 M to ensure alginate spheroid stability. Images were taken of the 24 well plates, and the images were analyzed as described in the section ‘Color analysis of graphene oxide reduction sample images’ to observe the reduction of graphene oxide over time. Following the measurement, spheroids were incubated at 30 °C with the media refreshed every two days to perform later measurements.

### Synthesis of CdSe quantum dots

Cadmium(II) acetate anhydrous (99.995%), selenium powder (100 mesh, 99.99%), tri-n-octyl phosphine (TOP, 97%), tri-n-octyl phosphine oxide (TOPO, 90%), hexadecylamine (HDA, 98%), tetradecyl phosphonic acid (TDPA, 97%), L-cysteine (L-Cys, 97%), and tetramethylammonium hydroxide pentahydrate (TMAHP, ≥97%) were acquired from Sigma Aldrich. HPLC-grade hexanes, diethyl ether, ethyl acetate, methanol, and acetone were acquired from Fischer Chemical.

CdSe quantum dots were synthesized following a hot-injection protocol published in Burke et al. (2021)^90^. Inside a nitrogen-filled glovebox, 0.79 g selenium powder was dissolved in 10 mL tri-n-octylphosphine (TOP). In a separate vial, 0.747 g anhydrous cadmium (II) acetate was dissolved in 12 mL TOP. The two solutions were stirred overnight at 50 °C for complete dissolution. Outside the glovebox, 7.8 g tri-n-octylphosphine oxide (TOPO), 2.3 g hexadecylamine (HDA), and 0.171 g tetradecylphosphonic acid (TDPA) were loaded into a 100-mL three-neck flask. The contents of the flask were purged with nitrogen gas and heated to 95 °C until a liquid viscosity was attained. The contents were degassed under a nitrogen pressure of under 0.1 Torr for 30 minutes. Afterwards, 1.8 mL of the TOP/Se solution was injected, and the temperature was increased to 315 °C. The temperature was then set to 260 °C, and 2.5 mL of the TOP/Cd solution was rapidly injected at 310 °C. The temperature was quickly adjusted to 260°C using an air gun. The reaction was maintained at 260 °C for 7 min, then cooled with an air gun and a water bath. Hexanes (15 mL) were added at 100 °C to prevent TOPO solidification. The crude product was split across two 50 mL Falcon tubes and precipitated with 10 mL of methanol and 25 mL of acetone in each tube, then centrifuged at 8000 rpm for 15 minutes. The pellet was redispersed in 20 mL of hexane and centrifuged at 8000 rpm for 15 minutes. The quantum dot solution was then collected in a scintillation vial, and the pink-white pellet was discarded.

### Ligand exchange to produce L-cysteine conjugated CdSe quantum dots

For ligand exchange reactions, 100 nM of CdSe quantum dots (QD) were calculated using the optical density at the first excitonic peak. The ligand to QD ratio was maintained at ∼1:6500 equivalents.

L-cysteine ligand exchange was performed guided by a previous procedure^63^. For 100 nM of QDs, a ligand solution was prepared by dissolving 0.242 g tetramethylammonium hydroxide pentahydrate (TMAH pentahydrate) and 0.116 g L-cysteine in 13.3 mL methanol to ensure deprotonation of thiol, amine, and carboxylate groups. A portion (8.3 mL) of this solution was placed in a 100-mL round-bottom flask under nitrogen gas, and 100 nM of freshly centrifuged (8000 rpm, 10 min) TDPA-CdSe QDs were loaded into a 1 mL syringe. The QDs were added dropwise to the ligand solution under stirring and active nitrogen gas flow, followed by the addition of ∼2 mL hexane. The mixture was refluxed at 50 °C for 45 minutes. This hot solution was precipitated by adding to ethyl ether (32.5 mL), followed by the addition of ethyl acetate (10 mL), and centrifuged at 4500 rpm for 15 min. The resulting pellet was dried under nitrogen and redispersed in Nanopure water to obtain L-cysteine-capped CdSe quantum dots.

### Hydrogen generation using cadmium selenide quantum dots as a catalyst

Hydrogen production experiments were conducted as outlined in Edwards et al. (2023)^42^. Prior to beginning photochemical experiments, 41-mL scintillation vials (Sigma-Aldrich) and custom gas-tight caps containing GC septa (Restek) were autoclaved. Each experimental solution was a total of 5.0 mL in modified minimal media (pH 7, see SI Appendix for additional details). After bacterial cultures were grown to an O.D._600_ of 0.4 to 0.5 (vide supra), approximately 500 µL of the cultures was transferred to minimal media to reach an O.D._600_ of 0.05 in photochemical vials. Where bacterial alginate structures were employed, these were used in place of overnight cultures and added in a 2:1 ratio of *ΔhydAΔhydB S. oneidensis-*containing alginate spheroids to *K. pneumoniae*-containing spheroids. CdSe–cysteine quantum dots were added to a final concentration of 1.0 µM, typically requiring about 230 µL from a CdSe–cysteine stock solution. Each 5 mL solution in a 41-mL vial was sealed with a gas-tight cap and a septum and deoxygenated by purging the solution for 20 min and headspace for 10 min with a 79.31%:20.69% N_2_:CH_4_ (Airgas) mixture using sterile needles. CH_4_ was present as a reference for product quantification. Illumination in the photochemical experiments was performed in a custom-built 16-sample apparatus into which 41-mL scintillation vials were placed. The samples were temperature controlled at 25 °C with a circulating water bath and maintained constant shaking at 100 rpm throughout the experiment. Samples were illuminated from below by a light-emitting diode (Philips LumiLED Luxeon Star Hex green 700 mA LEDs) at 530 nm (±10 nm). At the beginning of each experiment, the power of each LED was set to 25 mW ± 5 mW, as measured with a Nova II power meter (Ophir-Spiricon LLC) placed over each of the 16-wells. The amount of H_2_ produced was determined by sampling the headspace via gas chromatography (GC). To monitor H_2_ evolution, 25 μL of headspace gas was withdrawn from each vial through a septum with a 50 μL gas-tight syringe (Hamilton). Headspace gas samples were analyzed on a Shimadzu GC-2014AT gas chromatograph (GC) with a thermal conductivity detector and Carboxen 1010 PLOT column (30 m × 0.53 mm, Supelco) to quantify the H_2_ evolved with reference to CH_4_ (in the purging gas mixture) as the internal standard.

## Acknowledgements

*Shewanella oneidensis* knockout strains were kindly gifted by Dr. Benjamin Keith Keitz (UT Austin) and Dr. Jeffrey Gralnick (University of Minnesota). We gratefully acknowledge Dr. Allison J. Lopatkin (University of Rochester) for generously providing laboratory space and access to facilities, which enabled key bacterial experiments. We thank Pete Messmer at Lively Run Goat Dairy for the contribution of goat cheese waste runoff. This work was supported by the Chemical Sciences, Geosciences, and Biosciences Division, Office of Basic Energy Sciences, Office of Science, U.S. Department of Energy, Grant No. DE-SC0023354. Partial support for work by R. M. K. was from the National Institutes of Health, T32-GM118283.

