## Supplemental Figures 1-6 for "Transforming dairy waste into hydrogen fuel using alginate-encapsulated bacterial co-cultures"

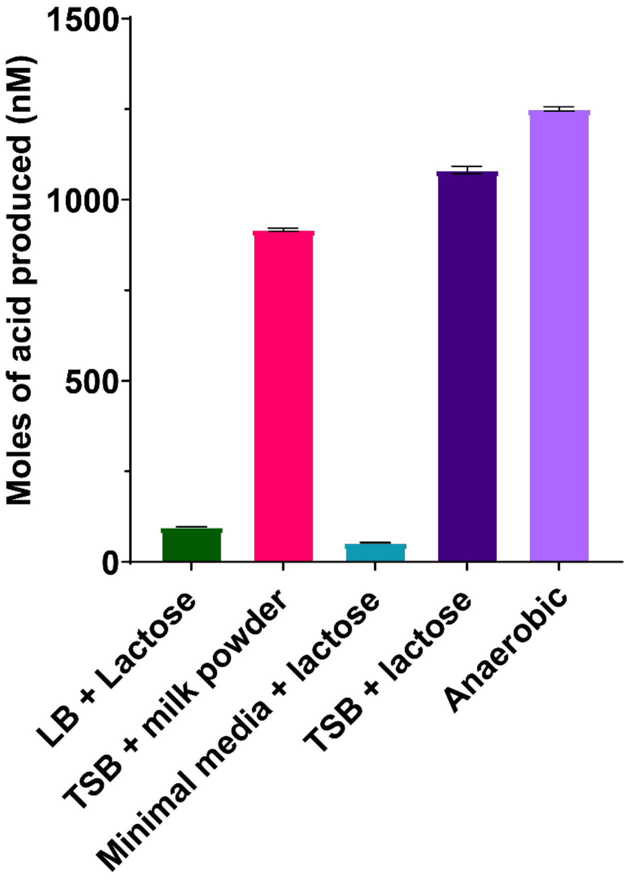


Supplemental Figure 1: Acid production of *Klebsiella pneumoniae* (KP). KP cultures were grown in various media at 30 ºC until a stable pH was reached. Acid production was calculated from the change in pH measured by a pH probe between initial and final pH readings, assuming total disassociation of the DL-lactic acid produced by the bacteria. All additions to TSB or LB media are present at 1% concentration in each sample well. Error bars represent the standard deviation of triplicate samples.

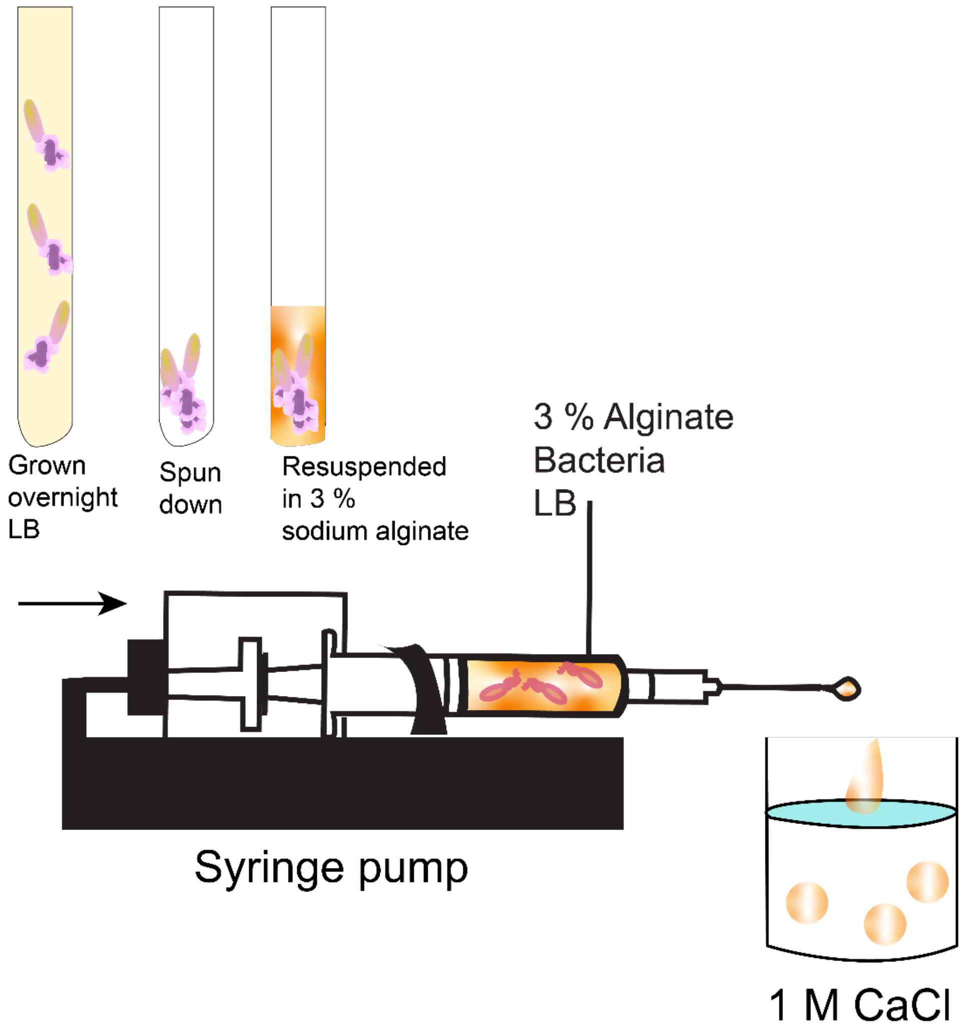


Supplemental Figure 2: The production of the alginate spheroids. Bacterial alginate bio-inks were prepared by spinning down overnight bacteria cultures and resuspending the bacterial pellets in 3% sodium alginate solution. The bio-ink was then extruded through a 10 mL syringe using a syringe pump into 0.1 M calcium chloride solution to form spheroids.

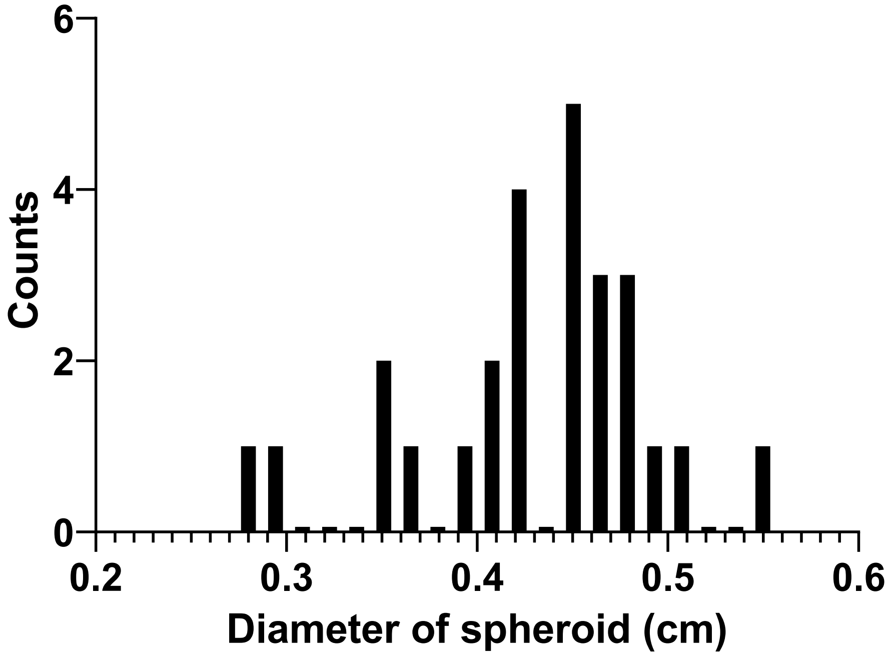


Supplemental Figure 3: Homogeneity of spheroids produced by syringe pump. Histogram shows the distribution of spheroid diameters. Spheroids were imaged from above, and diameters of individual spheroids were measured using ImageJ, using the diameter of the containing vessel was used as a reference.

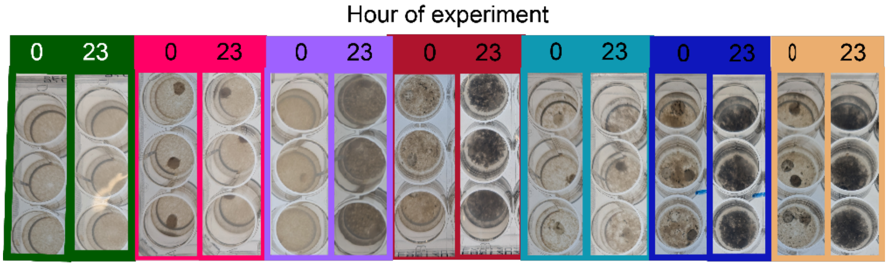


Supplemental Figure 4: Assays demonstrating improved extracellular reduction for co-cultures. Spheroids containing monocultures of *S. oneidensis* and *K. pneumoniae* were added to minimal media containing graphene oxide with added 20 mM lactic acid, 1% milk powder, or 1% whey. Images show the triplicate wells used to calculate the average HSV value change at 0 and 23 hours, demonstrating the value shift of the solutions that occurs with graphene oxide reduction. Outline colors of the wells correspond to the colors of the bars in Figure 4, namely: no bacteria (dark green), MR1 (pink), MR1 + milk (purple), KP + milk (burgundy), MR1 + 20 mM lactic acid (light blue), MR1 + KP + milk (dark blue), and MR1 + KP + whey (light orange).

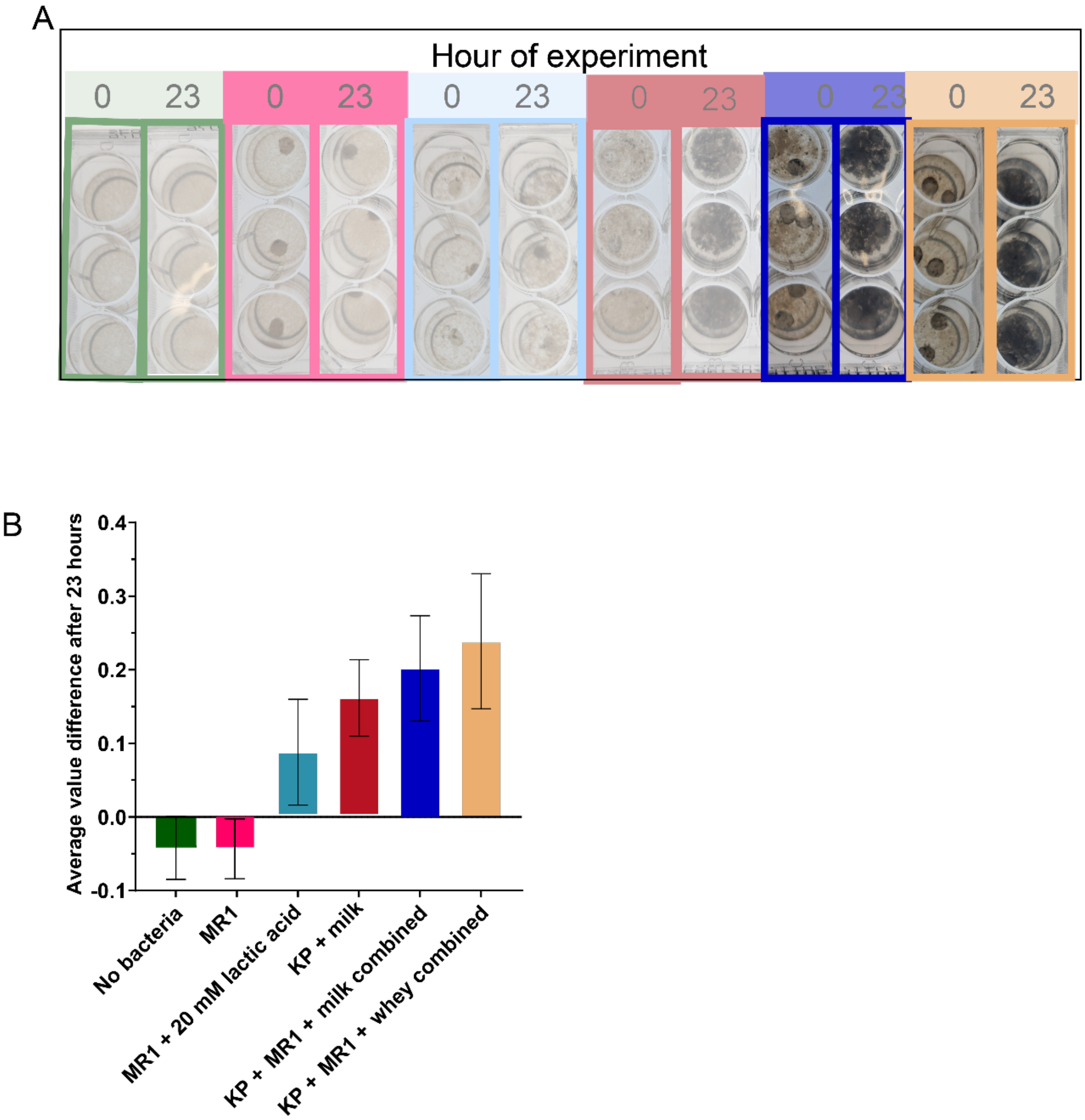


Supplemental Figure 5: Improvement in extracellular reduction from co-cultures when bacteria strains are combined within bio-ink spheroids. A) Spheroids containing monocultures or combined cultures of *S. oneidensis* and *K. pneumoniae* were added to minimal media containing graphene oxide with added 20 mM lactic acid, 1 %milk powder, or 1% whey. Spheroids were incubated in triplicate wells containing graphene oxide reduction reactions and imaged at 0 and 23 hours, and the average HSV value change was calculated for each condition. Media were refreshed throughout the experiment over 22 days. B) Cumulative average change in HSV value for graphene reduction reactions over 22 days of the experiment with each respective measurement representing a transferral of the same spheroids to fresh media. Bar colors correspond to the outline colors of the wells from panel A. Error bars represent the standard deviation of triplicate samples.

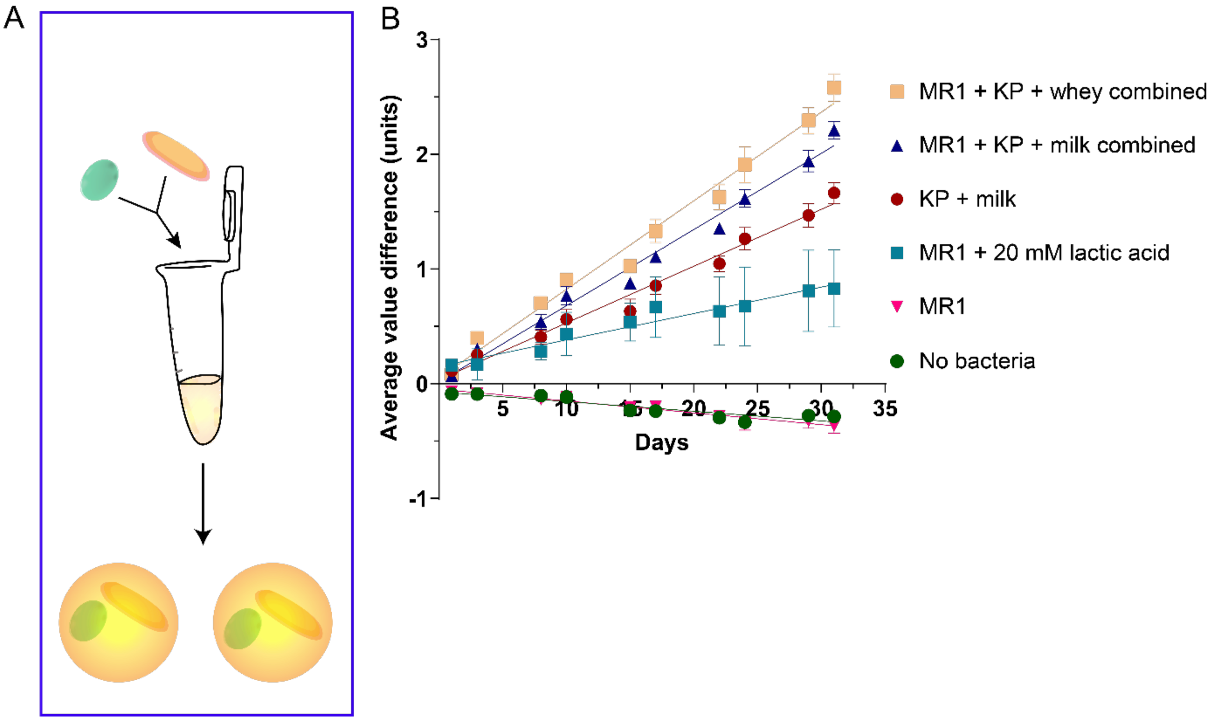


Supplemental Figure 6: Extended re-use of encapsulated co-cultures combined in bio-ink spheroids. A) *S. oneidensis* and *K. pneumoniae* cultures were combined and encapsulated within in the same alginate spheroids. B) Spheroids were added to minimal media containing 20 mM lactic acid, 1% milk powder or 1% whey. Images were taken at 0 hr and 23 hr timepoints, and the average change in HSV value of each sample was calculated. Media was refreshed for every following measurement. Cumulative average values are presented. Error bars represent the standard deviation of triplicate samples.
